# *Leishmania donovani* metacyclic promastigotes impair phagosome properties in inflammatory monocytes

**DOI:** 10.1101/2021.01.07.425828

**Authors:** Christine Matte, Guillermo Arango Duque, Albert Descoteaux

**Affiliations:** INRS – Centre Armand-Frappier Santé Biotechnologie, Université du Québec, Laval, QC, Canada

## Abstract

Leishmaniasis, a debilitating disease with clinical manifestations ranging from self-healing ulcers to life-threatening visceral pathologies, is caused by protozoan parasites of the *Leishmania* genus. These professional vacuolar pathogens are transmitted by infected sand flies to mammalian hosts as metacyclic promastigotes and are rapidly internalized by various phagocyte populations. Classical monocytes are among the first myeloid cells to migrate to infection sites. Recent evidence shows that recruitment of these cells contributes to parasite burden and to the establishment of chronic disease. However, the nature of *Leishmania*-inflammatory monocyte interactions during the early stages of host infection has not been well investigated. Here, we aimed to assess the impact of *Leishmania donovani* metacyclic promastigotes on antimicrobial responses within these cells. Our data showed that inflammatory monocytes were readily colonized by *L. donovani* metacyclic promastigotes, while infection with *Escherichia coli* was efficiently cleared. Upon internalization, metacyclic promastigotes inhibited superoxide production at the parasitophorous vacuole (PV) through a mechanism involving exclusion of NADPH oxidase subunits gp91^*phox*^ and p47^*phox*^ from the PV membrane. Moreover, we observed that unlike phagosomes enclosing zymosan particles, vacuoles containing parasites acidified poorly. Interestingly, whereas the parasite surface coat virulence glycolipid lipophosphoglycan was responsible for the inhibition of PV acidification, impairement of the NADPH oxidase assembly was independent of lipophosphoglycan and of the metalloprotease GP63. Collectively, these observations indicate that permissiveness of inflammatory monocytes to *L. donovani* may thus be related to the ability of this parasite to impair the microbicidal properties of phagosomes.

## Introduction

Monocytes are circulating mononuclear phagocytes that are critical orchestrators of the innate immune response to sterile and microbial inflammation (1, 2). They primarily originate from bone marrow myeloid progenitor cells and enter the circulation to infiltrate organs such as the liver and spleen. Monocytes can be classical (Ly6C^hi^) or non-classical (Ly6C^lo^), a dichotomy that allows them to originate distinct macrophage subsets (3, 4). Under physiological conditions, Ly6C^hi^ monocytes monitor extravascular tissues and present antigens to lymphocytes in lymph nodes. Inflammatory signals such as pathogen-derived molecules not only trigger myeloid differentiation into Ly6C^hi^ monocytes, but also their CCL2 chemokine-mediated migration to the injury site (5). There, these cells ingest microbes, release inflammatory molecules, and differentiate into macrophages and dendritic cells (1, 6). The importance of Ly6C^hi^ monocytes in antibacterial defense is channelled through their ability to phagocytose and kill pathogens (1, 6- 8). At the cellular level, phagosome maturation is driven by sequential soluble *N*-ethylmaleimide-sensitive factor attachment protein receptors (SNARE)-mediated interactions with endosomes and lysosomes (9, 10). This ensues in assembly of the NADPH and NOS2 oxidases on the phagosome membrane, which promote the intraluminal synthesis of the highly microbicidal reactive oxygen (ROS) and nitrogen (RNS) species, respectively (8, 10). Importantly, progressive acidification of the phagosome through the activity of the vacuolar ATPase (v-ATPase) promotes the function of pH-sensitive hydrolytic enzymes (11).

While Ly6C^hi^ monocytes mount a timely and effective response to injury, their persistent accrual may lead to excessive inflammation and organ damage (1). Similarly, Ly6C^hi^ monocytes have been reported to control (12, 13) or exacerbate (14-16) infection by protozoan parasites of the *Leishmania* genus. These parasites cause a group of anthropozoonoses called leishmaniases that are transmitted to mammalian hosts via the inoculation of metacyclic promastigotes by phlebotomine sand flies (17, 18). There, promastigotes are internalized into parasitophorous vacuoles (PV) by host phagocytes and differentiate into amastigotes which in turn propagate infection and disease. Studies in macrophages have revealed that metacyclic promastigotes build their intracellular niche using an armament of abundant GPI-anchored virulence-associated glycoconjugates such as the glycolipid lipophosphoglycan (LPG) and the metalloprotease GP63. These components of the promastigote surface coat were shown to alter phagosome maturation and inhibit acquisition of microbicidal properties, contributing to the colonization of host cells by *Leishmania* promastigotes (19-29).

When *L. major* or *L. amazonensis* are inoculated into the dermis, interferon gamma (IFN-γ)- and CCL2-mediated recruitment of Ly6C^hi^ monocytes creates a pool of cells that are readily infected by the parasites (15, 16, 30). Similarly, *L. donovani* parasites trigger the expansion of myeloid precursors, which augments the supply of Ly6C^hi^ monocytes to the viscera (14). The early events surrounding the phagocytosis of metacyclic promastigotes by inflammatory monocytes have not been elucidated. In this work, we observed that while inflammatory monocytes are efficient at clearing bacteria, they are highly permissive to *L. donovani* infection. We investigated the impact of *L. donovani* metacyclic promastigotes on inflammatory monocyte antimicrobial effector mechanisms and demonstrated that infection leads to an inhibition of PV oxidative activity and acidification.

## Results

### *L. donovani* metacyclic promastigotes successfully colonize inflammatory monocytes

Previous studies have provided evidence that Ly6C-positive inflammatory monocytes serve as permissive host cells for *L. major* during acute primary infection in the skin (30). To investigate how *Leishmania* colonize these cells, we assessed their permissiveness to *L. donovani* metacyclic promastigotes *in vitro*, as inflammatory monocytes also play a role in promoting parasitemia and susceptibility to visceral leishmaniasis (31, 32). We infected inflammatory monocytes generated from the bone marrow of C57BL/6 mice with serum-opsonized *L. donovani* metacyclic promastigotes and we assessed parasite burden over a period of 96 h. Consistent with their role as reservoirs that foster parasite burden (31, 32), inflammatory monocytes supported the proliferation of *L. donovani*, which doubled in number within 96 h (Fig. 1A). Classical and intermediate monocytes, compared to their non-classical counterparts, possess efficient antimicrobial effector mechanisms, including powerful oxidative capacities exerted via NADPH oxidase activity (33-35). In murine infection models, their ability to restrict the growth of pathogens such as *Mycobacterium tuberculosis, Salmonella*, or *Listeria* has been well established (8, 35-37). To rule out any potential general defect in the microbicidal response of *in vitro*-generated inflammatory monocytes, we fed them live *Escherichia coli* and assessed bacteria killing over time. In contrast to *L. donovani* metacyclic promastigotes, bacteria were rapidly cleared within 4 h (Fig. 1B). Pretreatment of inflammatory monocyte cultures with the NADPH oxidase inhibitor diphenyleneiodonium (DPI) significantly reduced bactericidal activity compared to vehicle drug alone (DMSO), confirming the contribution of reactive oxygen species (ROS) in the elimination of internalized microbes by these cells (Fig. 1C).

**Figure 1.**
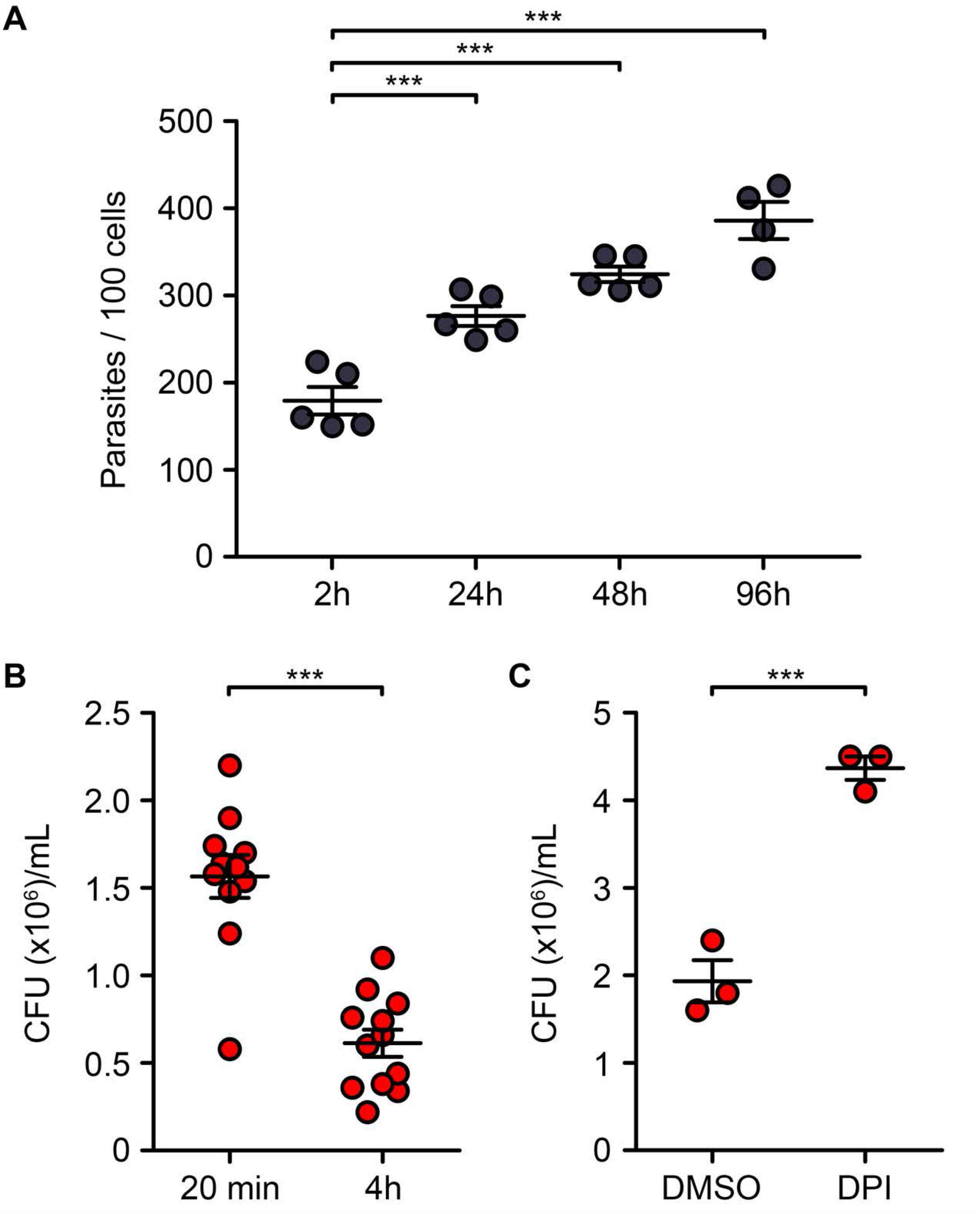
Inflammatory monocytes are permissive to *L. donovani* metacyclic promastigotes replication. Bone marrow cells from C57BL/6 mice were incubated for 3 days in the presence of M-CSF. Non-adherent cells were seeded in chambers mounted on Permanox slides and stimulated for 3 h with 100 U/mL IFN-γ. Cells were then incubated with serum-opsonized, *L. donovani* LV9 metacyclic promastigotes. After 2 h, non-attached/non-internalized parasites were removed by washing and intracellular parasitemia was assessed at the indicated time points by counting the number of parasites per 100 cells, upon staining with the Hema 3 staining kit **(A)**. Inflammatory monocytes were left untreated **(B)** or treated for 1 h with the NADPH oxidase inhibitor DPI (10 μM) or vehicle drug (DMSO 0.1%) **(C)** and then infected with bacteria (*E. coli* DH1; OD_660nm_ *=* 0.6; bacteria-to-cell ratio of 20:1) for 20 min. Non-internalized bacteria were removed by washing and via incubation in the presence of 5 μg/mL gentamycin for 20 min **(B, C)** or 4 h **(B)**, after which the amount of viable intracellular bacteria was assessed through lysis, serial dilutions and counting of CFUs on agar plates. The means ± SEM of data from two independent experiments representative of a total of four **(A)**, from four independent experiments **(B)** or from one experiment representative of two **(C)** are shown. *** *P* < 0.001 according to a one-way ANOVA with Bonferroni *post-hoc* test **(A)** or to a two-tailed, unpaired t-test **(B, C)**.

### *L. donovani* promastigotes prevent assembly of the NADPH oxidase at the PV in inflammatory monocytes

Given the role of ROS in the antimicrobial activity of inflammatory monocytes, we examined the possibility that *L. donovani* metacyclic promastigotes colonize those cells by interfering with the production of ROS. Inflammatory monocytes were fed opsonized parasites for 15 min and then incubated with NitroBlue Tetrazolium (NBT), a compound that generates dark formazan deposits upon reduction by superoxide. After 1 h, we detected formazan in 30% of PVs harboring *L. donovani* (Fig. 2). On the other hand, internalization of zymosan led to the generation of formazan in over 65% of phagosomes (Fig. 2). These results indicated that *L. donovani* metacyclic promastigotes impair phagosomal ROS production in inflammatory monocytes. To identify the mechanism underlying the impaired phagosomal ROS production in *L. donovani*-infected inflammatory monocytes, we investigated the assembly of the NADPH oxidase at the PV membrane. The cytosolic component p47^*phox*^ and the transmembrane subunit gp91^*phox*^ are integral in mediating NADPH oxidase assembly and function, respectively. While p47^*phox*^ orchestrates complex formation in response to activation by translocating other cytosolic subunits to the membrane, gp91^*phox*^ catalyzes superoxide production by transferring electrons from NADPH to molecular oxygen (38). Using confocal immunofluorescence microscopy, we monitored the redistribution of each subunit in Ly6C-positive monocytes upon phagocytosis of zymosan *versus* promastigotes. At 15 and 60 min post-internalization, phagosomes containing zymosan displayed evident recruitment of gp91^*phox*^ and p47^*phox*^ (Fig. 3). Interestingly, Ly6C was also detected on the phagosome membrane, possibly as a result of plasmalemma internalization during formation and sealing of the phagocytic cup. Conversely, the NADPH oxidase subunits were excluded from the majority of *L. donovani*-harboring PVs, indicating that metacyclic promastigotes interfere with assembly of the NADPH oxidase to quell ROS production in inflammatory monocytes.

**Figure 2.**
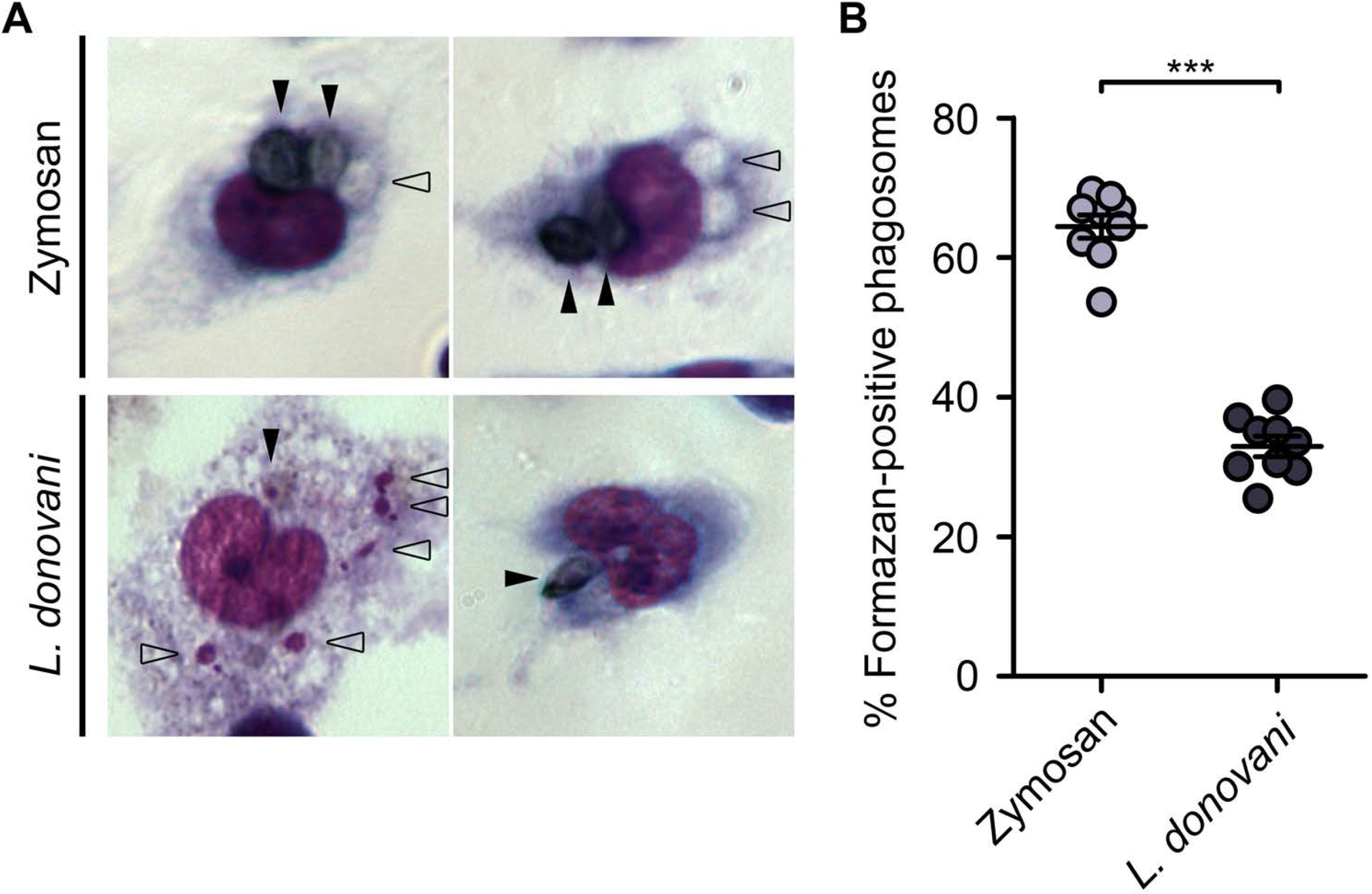
*L. donovani* metacyclic promastigotes inhibit phagosomal ROS production. Inflammatory monocytes were incubated for 15 min with serum-opsonized zymosan particles or *L. donovani* LV9 metacyclic promastigotes, after which non-attached/non-internalized particles were removed by washing. Cells were incubated for 1 h with 1 mg/mL NBT and then stained with the Hema 3 staining kit. Phagosomes positive (full arrowheads) or negative (empty arrowheads) for the presence of formazan deposits were counted. Representative images **(A)** and the means ± SEM of data from three independent experiments **(B)** are shown. *** *P* < 0.001 according to a two-tailed, unpaired t-test.

**Figure 3.**
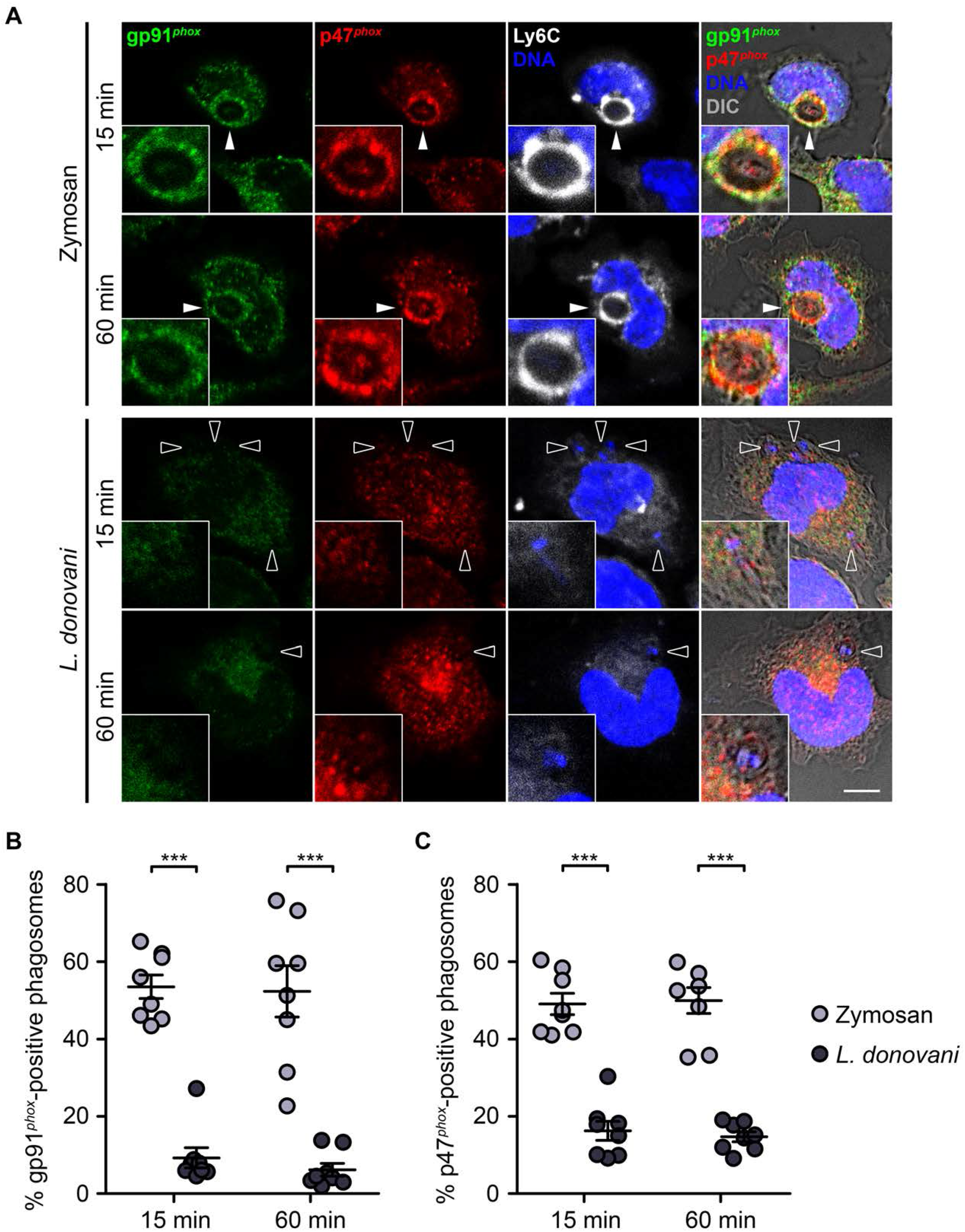
*L. donovani* metacyclic promastigotes inhibit phagosomal NADPH oxidase assembly. Inflammatory monocytes were incubated for 15 min with serum-opsonized zymosan particles or *L. donovani* WT metacyclic promastigotes, after which non-attached/non-internalized particles were removed by washing. Cells were further incubated for the indicated time points before being fixed, blocked, permeabilized and stained with DAPI (DNA, blue) and with antibodies against gp91^*phox*^ (green), p47^*phox*^ (red) and Ly6C (white). Ly6C-positive monocytes were then visualized by confocal immunofluorescence microscopy to quantify zymosan- and promastigote-containing phagosomes that were positive (full arrowheads) or negative (empty arrowheads) for gp91^*phox*^ or p47^*phox*^ recruitment to the membrane. Representative images **(A)** and the means ± SEM of data from three independent experiments **(B, C)** are shown. Scale bar, 5 μm. DIC: differential interference contrast. *** *P* < 0.001 according to a one-way ANOVA with Bonferroni *post-hoc* test.

### *L. donovani* promastigotes impairs PV acidification

Phagosomal acidification is required for the intracellular elimination of microorganisms by macrophages, since lysosomal enzymes are optimally active at acidic pH (39). We previously reported that *L. donovani* and *L. braziliensis* promastigotes inhibit phagosome acidification during the early phases of macrophage infection (40, 41). Hence, we next assessed the potential impact of *L. donovani* on PV acidification in Ly6C-positive inflammatory monocytes using the acidotropic probe LysoTracker DND-99 and confocal microscopy. We used zymosan as a positive control for phagosome acidification. After 2 h, most phagosomes containing zymosan were fluorescent due to protonation of the probe, indicative of an acidic lumen, and the percentage continued to increase over time (Fig. 4). Of note, a general accumulation of acidic vesicles was observed in the cytosol of Ly6C-positive monocytes at 6 h post-phagocytosis of zymosan (Fig. 4A). In contrast, *L. donovani* metacyclic promastigotes were enclosed in PVs that were predominantly negative for LysoTracker fluorescence at 2 h and 6 h post-infection (Fig. 4), indicating that these parasites inhibit PV acidification in Ly6C-positive monocytes.

**Figure 4.**
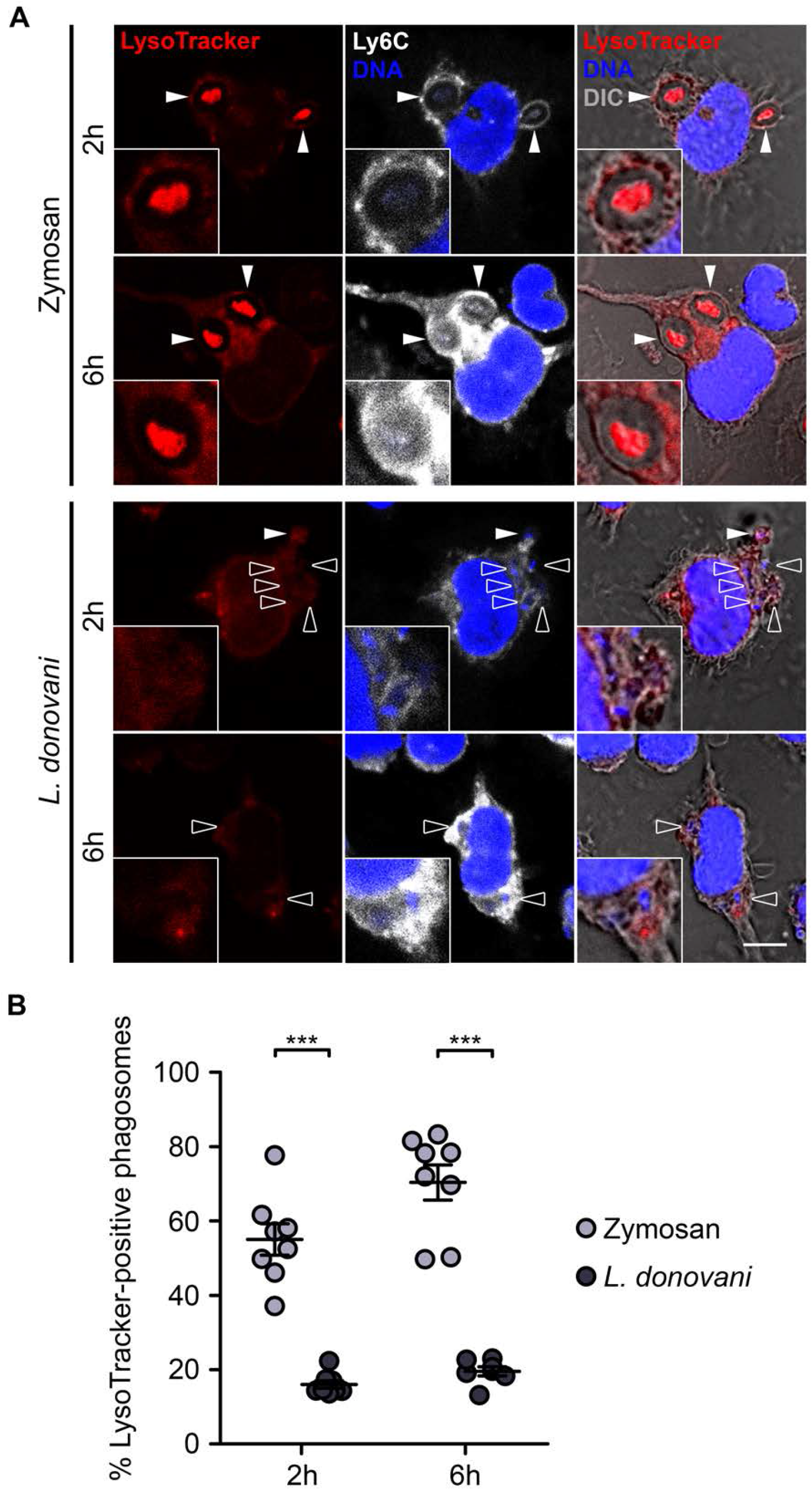
*L. donovani* metacyclic promastigotes inhibit phagosomal acidification. Inflammatory monocytes were incubated for 15 min with serum-opsonized zymosan particles or *L. donovani* WT metacyclic promastigotes, after which non-attached/non-internalized particles were removed by washing. Cells were further incubated for up to 6 h in the presence of LysoTracker DND-99 (red), added 2 h before the end of each time point. Cells were then fixed, blocked, permeabilized and stained with DAPI (blue) and for Ly6C (white). Ly6C-positive monocytes were subsequently visualized by confocal immunofluorescence microscopy to quantify the number of phagosomes that were positive (full arrowheads) or negative (empty arrowheads) for LysoTracker fluorescence. Representative images of the 2 h time point **(A)** and the means ± SEM of data from three independent experiments **(B)** are shown. Scale bar, 5 μm. DIC: differential interference contrast. *** *P* < 0.001 according to a one-way ANOVA with Bonferroni *post-hoc* test.

### LPG prevents PV acidification but not NADPH oxidase assembly in inflammatory monocytes

We previously reported that LPG and the metalloprotease GP63, two components of the *Leishmania* promastigote surface coat (42, 43), inhibit phagosomal assembly of the NADPH oxidase in macrophages through distinct mechanisms (23, 26). To determine the potential role of these virulence factors in the inhibition of phagosomal NADPH oxidase assembly in inflammatory monocytes, we first infected these cells with WT, LPG-deficient Δ*lpg1*, or phosphoglycan (Gal(*β*1,4)Man(*α*1)-PO_4_ repeating unit)-deficient Δ*lpg*2 *L. donovani* metacyclic promastigotes (Fig. S1). Inflammatory monocytes fed with zymosan were used as positive controls. At 1 h post-phagocytosis, we examined the recruitment of gp91^*phox*^ (Fig. 5A) and p47^*phox*^ (Fig. 5B) to the PV membrane of Ly6C-positive monocytes. Up to 60% of phagosomes containing zymosan were positive for gp91^*phox*^ and p47^*phox*^. In contrast, only 10 to 15% phagosomes harboring either WT, Δ*lpg1*, or Δ*lpg*2 parasites were positive for gp91^*phox*^ and p47^*phox*^. In agreement with these observations, we found that Δ*lpg1* and Δ*lpg2 L. donovani* metacyclic promastigotes dampened the production of superoxide in PVs to the same extent as WT parasites (Fig. 5C). We next assessed the potential role of GP63 in the inhibition of phagosomal NADPH oxidase assembly in inflammatory monocytes. Since no GP63-deficient parasites are available in *L. donovani*, we used WT, Δ*gp63* and Δ*gp63*+*GP63 L. major* metacyclic promastigotes to infect inflammatory monocytes. As shown in Figure 6, at 30 min post-phagocytosis, between 15 to 20% of phagosomes harboring these three *L. major* lines were positive for gp91^*phox*^ compared to 50% of phagosomes containing zymosan. Collectively, these results indicate that neither Gal(*β*1,4)Man(*α*1)-PO_4_-containing glycoconjugates nor GP63 are individually responsible for the inhibition of phagosomal NADPH oxidase assembly in inflammatory monocytes. Of note, the capacity of LPG- and GP63-defective metacyclic promastigotes to prevent phagosomal assembly of the NADPH oxidase is however not sufficient to survive in inflammatory monocytes (Fig. S2), consistent with the impact of these two virulence factors on multiple host cell pathways and processes (44-46).

**Figure 5.**
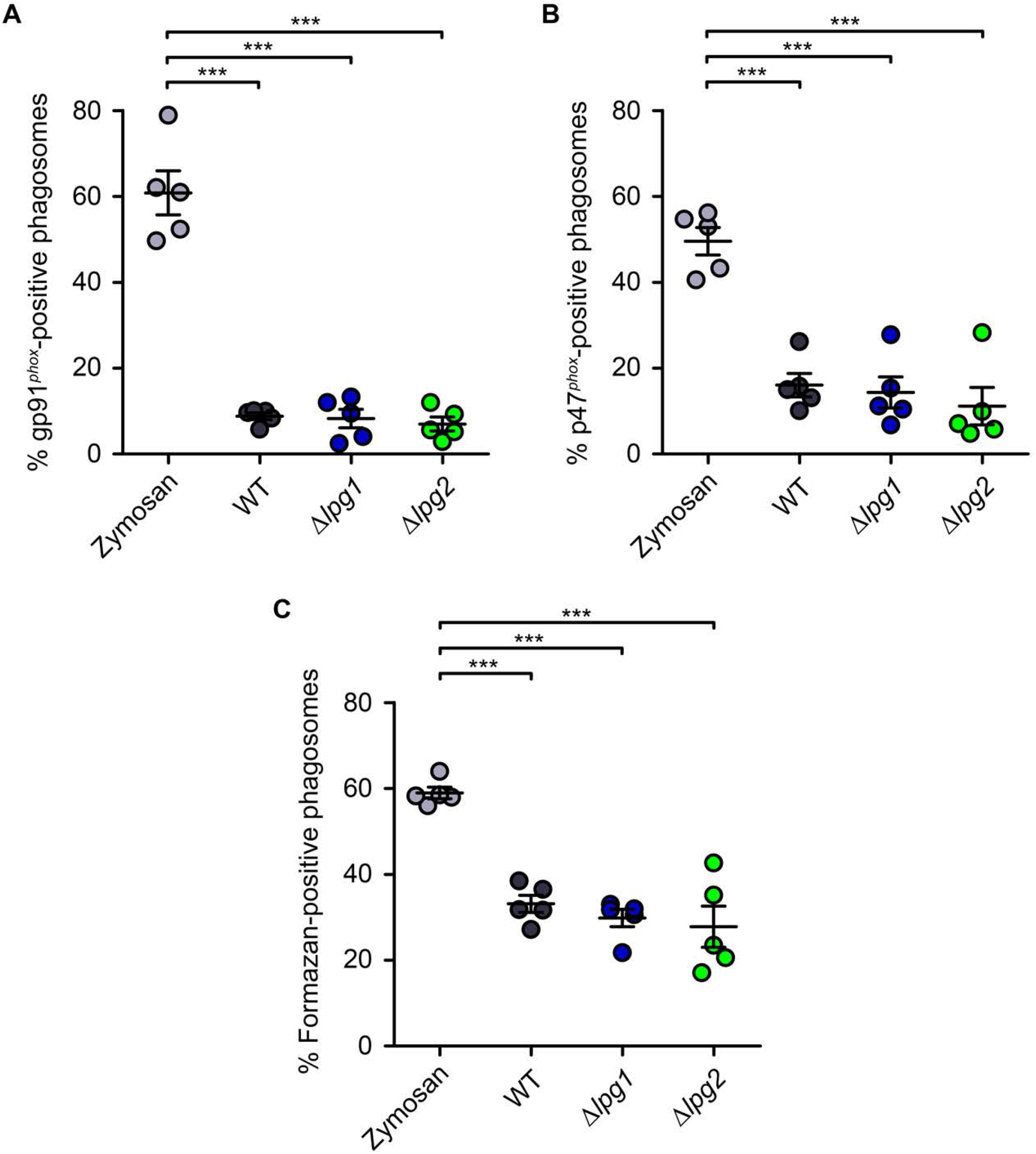
Inhibition of phagosomal ROS production is independent of phosphoglycans. Inflammatory monocytes were incubated for 15 min with serum-opsonized zymosan particles or *L. donovani* metacyclic promastigotes (WT, Δ*lpg1* and Δ*lpg2*), after which non-attached/non-internalized particles were removed by washing **(A-C)**. Cells were then further incubated for 1 h before being fixed, blocked, permeabilized and stained with DAPI (blue) and antibodies against gp91^*phox*^ (green), p47^*phox*^ (red) and Ly6C (white). Ly6C-positive monocytes were visualized by confocal immunofluorescence microscopy **(A, B)**. Alternatively, cells were incubated for 1 h with 1 mg/mL NBT prior to staining with the Hema 3 kit and visualization by light microscopy **(C)**. Phagosomes were evaluated for gp91^*phox*^ recruitment **(A)**, p47^*phox*^ recruitment **(B)** or the presence of formazan deposits **(C)**. The means ± SEM of data from three **(A, B)** or two **(C)** independent experiments are shown. *** *P* < 0.001 according to a one-way ANOVA with Bonferroni *post-hoc* test.

**Figure 6.**
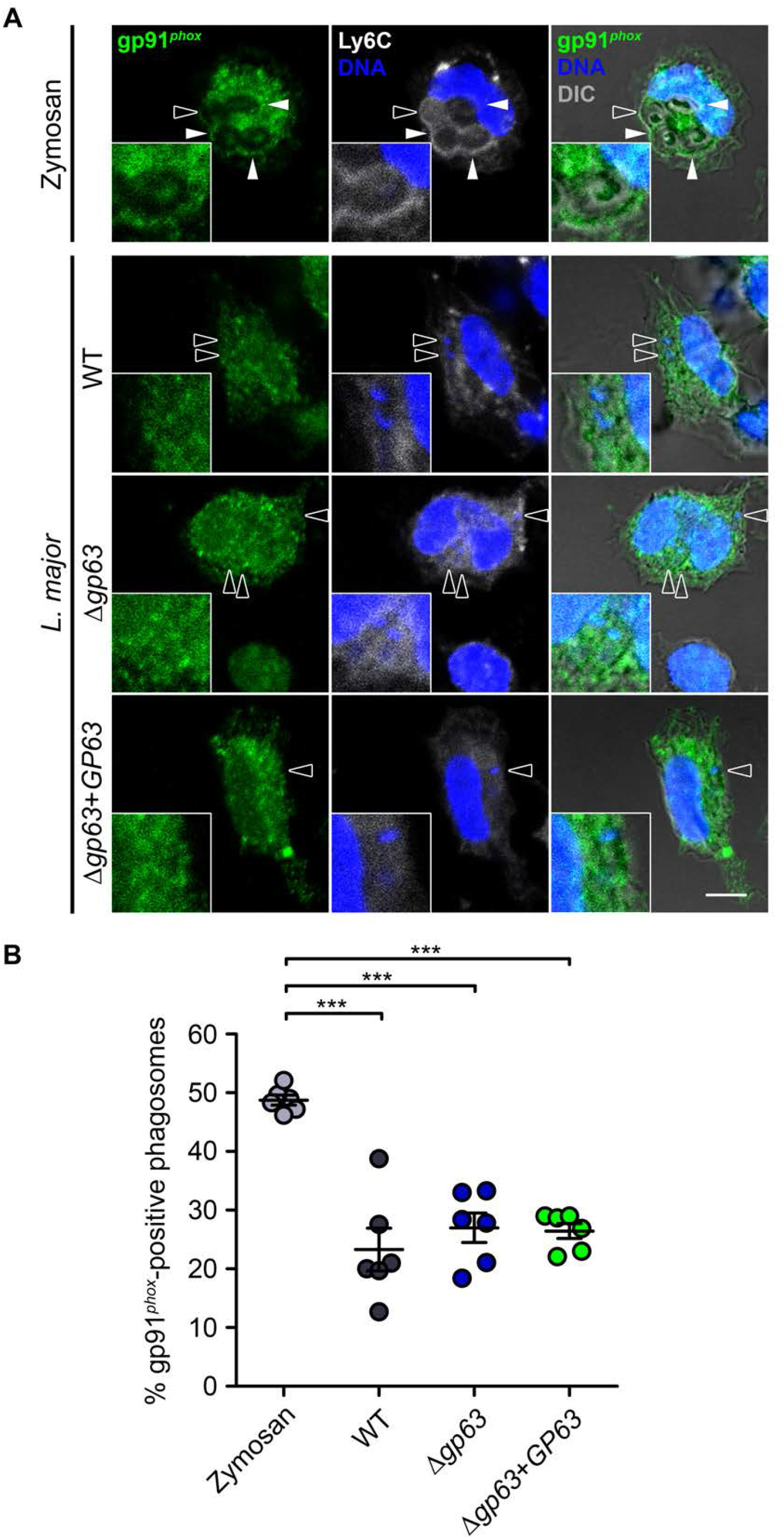
Inhibition of the phagosomal recruitment of gp91^*phox*^ is independent of GP63. Inflammatory monocytes were incubated for 15 min with serum-opsonized zymosan particles or *L. major* NIH S clone A2 promastigotes (WT, Δ*gp63* or Δ*gp63*+*GP63*), at particle-to-cell ratios of 3:1 and 10:1 respectively, after which non-attached/non-internalized particles were removed by washing. Cells were further incubated for 30 min before being fixed, blocked, permeabilized and stained with DAPI (DNA, blue) and with antibodies against gp91^*phox*^ (green) and Ly6C (white). Ly6C-positive monocytes were then visualized by confocal immunofluorescence microscopy to quantify zymosan- and promastigote-containing phagosomes that were positive (full arrowheads) or negative (empty arrowheads) for gp91^*phox*^ recruitment to the membrane. Representative images **(A)** and the means ± SEM of data from three independent experiments **(B)** are shown. Scale bar, 5 μm. DIC: differential interference contrast. *** *P* < 0.001 according to a one-way ANOVA with Bonferroni *post-hoc* test.

We previously reported that, in macrophages, LPG mediates exclusion of the v-ATPase from *L. donovani*-containing phagosomes (22, 24, 40). Using the Δ*lpg1* and Δ*lpg2* mutants, we verified the role of LPG and other Gal(*β*1,4)Man(*α*1)-PO_4_-containing parasite glycoconjugates in the inhibition of phagosome acidification in Ly6C-positive monocytes. Inflammatory monocytes fed with zymosan were used as positive controls. As depicted in Figure 7, in contrast to WT metacyclic promastigotes, the majority of Δ*lpg1* and *Δlpg2* parasites were present in acidic vacuoles. Finally, complementation of knockout strains (Δ*lpg1*+*LPG1* and Δ*lpg2*+*LPG2*) rescued the capacity of metacyclic promastigotes to prevent PV acidification (Fig. 7). Collectively, these results demonstrate that *L. donovani* metacyclic promastigotes prevent acidification of inflammatory monocyte phagosomes in a LPG-dependent manner. The absence of other PGs in Δ*lpg2* mutants did not affect the frequency of fluorescent PVs.

**Figure 7.**
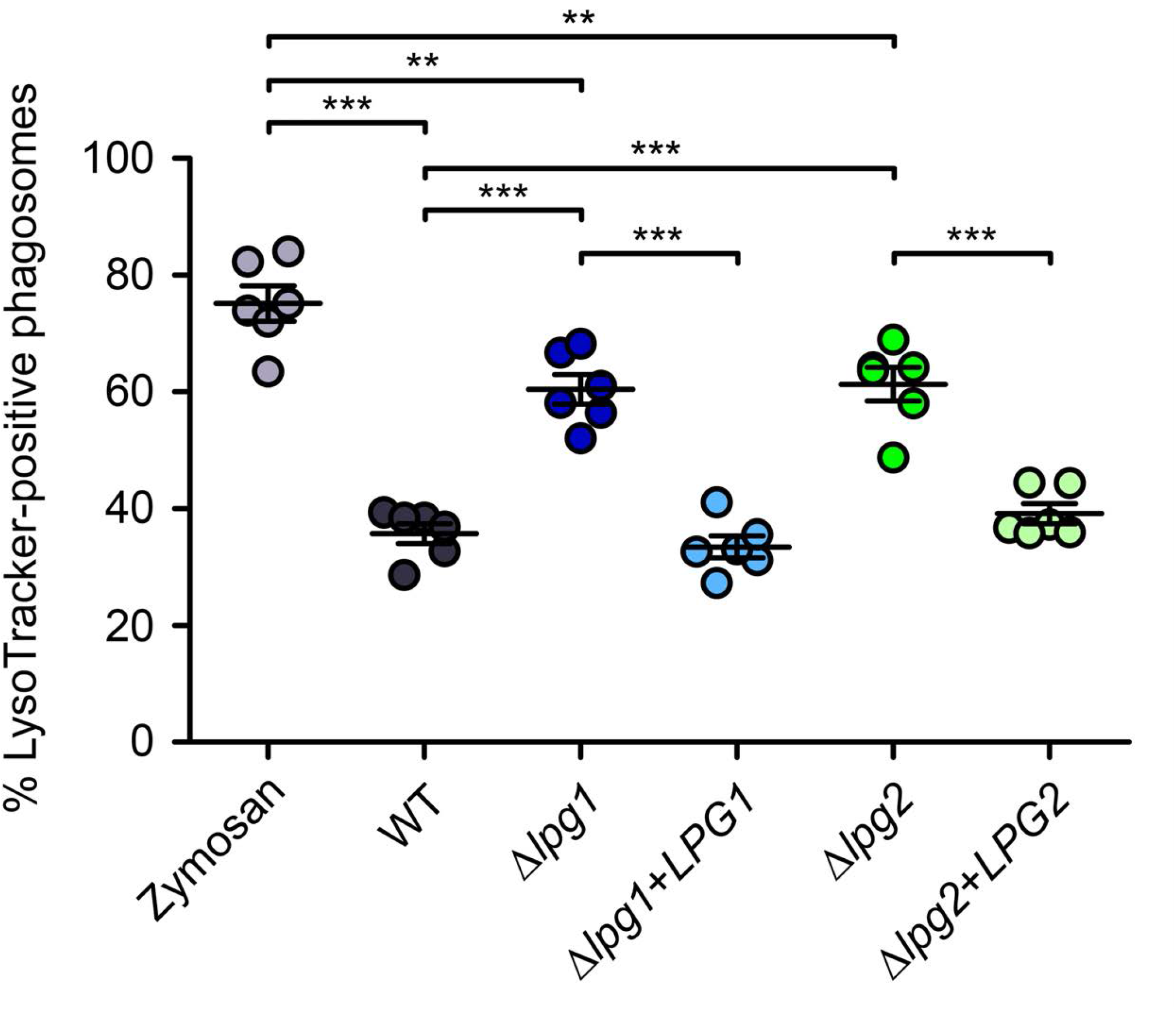
LPG is responsible for the inhibition of phagosomal acidification in inflammatory monocytes. Inflammatory monocytes were incubated for 15 min with serum-opsonized zymosan particles or *L. donovani* metacyclic promastigotes (WT, Δ*lpg1*, Δ*lpg2*, Δ*lpg1*+*LPG1*, Δ*lpg2*+*LPG2*), after which non-attached/non-internalized particles were removed by washing. Cells were then further incubated for 2 h in the presence of LysoTracker DND-99 (red) before being fixed, blocked, permeabilized and stained with DAPI (blue) and an antibody against Ly6C (white). Ly6C-positive monocytes were finally visualized by confocal immunofluorescence microscopy to assess phagosome acidification. The means ± SEM of data from three independent experiments are shown. *** *P* < 0.001 according to a one-way ANOVA with Bonferroni *post-hoc* test.

## Discussion

*Leishmania* metacyclic promastigotes are deposited by infected sand flies into the dermis of mammals where they are internalized by various phagocyte populations (30, 47-49). These include inflammatory monocytes which, despite their potent antimicrobial capacities, are highly permissive to *L. major* replication (30). In this study, we aimed to characterize the events that unfold upon phagocytosis of *L. donovani* metacyclic promastigotes by these cells. Our main finding was that *L. donovani* abrogate key antimicrobial processes, namely phagosomal assembly of the NADPH oxidase and acidification, which may contribute to their capacity to proliferate in inflammatory monocytes. The differential roles of parasite surface glycoconjugates in interfering with these inflammatory monocyte antimicrobial mechanisms are also described.

The outcome of pathogen uptake by phagocytic cells largely depends on the efficiency of microbicidal effector mechanisms within the newly formed phagosome. Generation of ROS at the phagosome represents one of the central host defense mechanism against infection and depends on the assembly of a functional NADPH oxidase (50, 51). To avoid the harmful consequences of exposure to ROS, pathogens have evolved a panoply of strategies to block assembly and/or resist activity of the NADPH oxidase (50). Hence, *Leishmania* uses diverse mechanisms to reduce exposure to ROS within mammals. One such mechanism is the conversion of toxic ROS through the action of enzymes such as superoxide dismutases (52, 53) and tryparedoxin peroxidase (54, 55). *Leishmania* also interferes with the production of ROS. Induction of higher expression of the macrophage heme oxygenase-1 (56-58) increases heme degradation and was shown to prevent gp91^*phox*^ maturation (56). Using LPG- and GP63-deficient mutants, we previously reported that in macrophages, disruption of phagosomal membrane lipid microdomains by LPG interferes with the recruitment of cytosolic p47*/*p67/p40^*phox*^ hetero-trimers (23), whereas recruitment of gp91^*phox*^ was rather shown to be inhibited in a GP63-dependent manner (29). Our observation that LPG- and GP63-defective mutants effectively prevented phagosomal assembly of the NADPH oxidase and ROS production to the same extent as WT promastigotes in inflammatory monocytes was surprising. Variations between the experimental system of the work presented here and that of previous studies, such as strain differences or the use metacyclic promastigotes rather than stationary-phase cultures, could also explain the discrepancies in the reported interplay between *Leishmania* virulence factors and the host cell oxidative response. Hence, we cannot exclude that with metacyclic promastigotes, the absence of LPG in our *L. donovani* Δ*lpg1* and Δ*lpg2* mutants was compensated by the action of GP63 on the NADPH oxidase. Conversely, LPG may have enabled *L. major* Δ*gp63* parasites with the capacity to block phagosomal assembly of the NADPH oxidase. Alternatively, there may be a cell type-specific divergence of pathways linking phagocytosis to NADPH oxidase assembly (59). The subcellular distribution of the transmembrane subunit gp91^*phox*^, for instance, varies between cell lineages. In resting neutrophils, gp91^*phox*^ resides in an intracellular pool of granules that are rapidly mobilized to the plasma membrane upon phagocytosis or other stimuli (60, 61). In dendritic cells and macrophages, gp91^*phox*^ localizes to both the plasma membrane and the endocytic recycling compartment and is replenished through fusion with vesicles of the endocytic compartment, via SNAREs such as VAMP8 (26, 61-63). Endothelial cells, on the other hand, maintain a reservoir of gp91^*phox*^ in the vicinity of the endoplasmic reticulum (64). In inflammatory monocytes, how membrane-bound NADPH oxidase subunits traffic to the phagosome has not been addressed. Nonetheless, it is possible that *Leishmania* parasites have evolved complementary strategies to subvert phagosome oxidative activity based on the biological processes of their various host cells. In this regard, *Leishmania* is known to trigger SHP-1 activation (65, 66), which was shown to inhibit phagosomal activation of NOX2 through the dephosphorylation of p47^*phox*^ (67). Whether such a mechanism is operational in infected inflammatory monocytes will deserve further investigation. Interestingly, Spath *et al*. previously reported that, in macrophages from gp91^*phox*^-deficient mice, a *L. major* Δ*lpg1* mutant survived as well as wild type parasites during the first two days but were unable to survive at later time points (21). These observations are consistent with a role for LPG in the ability of *Leishmania* promastigotes to impair and resist to the production of ROS and point to a broader role for LPG in the establishment of the parasite within host macrophages.

Phagosomes in classical monocytes and M1 macrophages have been reported to maintain an alkaline or near-neutral pH for up to 30 min post-phagocytosis, due to consumption of lumen protons by the highly active NADPH oxidase (33, 59). However, in both cell types, phagosome maturation eventually involves progressive acidification (10). The insertion of LPG into cell membrane bilayers is known to alter phagosome fusion with compartments of the endocytic pathway, leading to a transient arrest in maturation (19, 22, 68). While *L. donovani* promastigote-induced PVs acquire some characteristics of mature phagolysosomes at approximately 5 h post-infection, such as LAMP1 recruitment (69), the exclusion of the proton pump v-ATPase persists up to 24 h (40). This phenotype is congruent to what we observed here in inflammatory monocytes infected with *L. donovani* metacyclic promastigotes, confirming the premise that LPG-mediated inhibition of PV acidification is part of the colonization process of these cells. The observed impairment of PV acidification during the early stages of phagocyte colonization by *L. donovani* and *L. braziliensis* in both inflammatory monocytes and macrophages (40, 41) raises questions on the current model pertaining to the environmental cues associated to the differentiation of promastigotes into amastigotes. Indeed, it is assumed that transformation of promastigotes into amastigotes within the phagosome is mainly triggered by a rapid and concomitant exposure to an acidic environment and to an elevated temperature (70, 71). Our data challenge this model which has been widely used in *in vitro* studies aimed at elucidating the molecular events associated to the promastigote-to-amastigote differentiation process inside phagocytic cells. Clearly, additional studies will be required to elucidate this crucial aspect of *Leishmania* biology.

Prior to this study, the functional biology of inflammatory monocytes in the context of a primary infection with *L. donovani* metacyclic promastigotes had not been well addressed. The data presented here demonstrate that, like macrophages, inflammatory monocytes fail to generate robust ROS production upon internalization of *L. donovani* and display a defect in phagosome acidification. Collectively, these findings provide us with a better insight into how inflammatory monocytes foster parasite proliferation and are consistent with the role of these host cells in the establishment of persistent visceral disease (31).

## Materials and methods

### Ethics statement

Mice were handled in strict accordance with good animal practice as defined by the Canadian Council on Animal Care. Experimental procedures were approved by the Comité Institutionnel de Protection des Animaux of the Centre National de Biologie Expérimentale (Protocol 1706-07).

### Monocyte cultures

To obtain bone marrow-derived monocytes, bone marrow was flushed from the femurs and tibias of 6- to 8-week old C57BL/6 mice, treated with 0.17 M NH_4_Cl pH 7.4 for 7 min to lyse red blood cells, and cultured in tissue culture-treated petri dishes in complete medium (Dulbecco’s Modified Eagle Medium with high glucose and L-glutamine (Life Technologies) containing 10% heat-inactivated fetal bovine serum (FBS) (Sigma-Aldrich), 10 mM HEPES pH 7.4 and Penicillin-Streptomycin (Life Technologies)) supplemented with 15% (v/v) L929 cell-conditioned medium (LCM) as a source of colony-stimulating factor-1 (CSF-1), in a 37°C incubator with 5% CO_2_. After three days, non-adherent cells were collected, transferred to Nunc Lab-Tek 8-well chambers fixed on Permanox plastic slides (Thermo Scientific) in complete medium without LCM and stimulated with 100 U/mL IFN-γ for 3h. The resulting inflammatory monocyte populations were verified by confocal immunofluorescence microscopy (see below) for Ly6C positivity and for the absence of polymorphonuclear cells (Fig. S4).

### Parasites cultures

*L. donovani* LV9 (MHOM/ET/67/HU3) and *L. major* NIH S (MHOM/SN/74/Seidman, clone A2 promastigotes were cultured at 26°C in *Leishmania* medium (M199 medium (Sigma Aldrich) supplemented with 10% heat-inactivated FBS, 10 mM HEPES pH 7.4, 100 μM hypoxanthine, 5 μM hemin, 3 μM 6-biopterin, 1 μM biotin and penicillin-streptomycin). The *L. donovani* LV9 isogenic Δ*lpg1* (LPG-defective) and Δ*lpg2* (PG-defective) mutants, produced via homologous gene replacement using targeting constructions previously described (72, 73), were cultured in the presence of hygromycin B, at the respective concentrations of 100 μg/mL and 300 μg/mL. To restore *LPG1* expression, Δ*lpg1* promastigotes were electroporated with pLeishZeo-*LPG1* (74) and complemented parasites (Δ*lpg1*+*LPG1*) were selected with 100 μg/mL zeocin (along with hygromycin B). To restore *LPG2* expression, Δ*lpg2* mutants were electroporated with pLeishNeo-*LPG2* (75) and complemented parasites (Δ*lpg2*+*LPG2*) were selected with 20 μg/mL G418 (in addition to hygromycin B). These parasite lines were regularly verified by agglutination tests and by Western blotting using the phosphoglycan-specific CA7AE antibody (MediMabs) (Fig. S1) (74). *L. major* NIH S clone A2 isogenic Δ*gp63* mutants and their complemented counterparts (Δ*gp63*+*GP63*, cultured in the presence of 50 μg/ml G418) have been previously described (76). GP63 expression was routinely assessed by Western blotting using the monoclonal antibody #235 provided by Dr. W. Robert McMaster (data not shown) (77). For infections, metacyclic promastigotes were enriched from late stationary-phase cultures using Ficoll gradients, as described before (78, 79).

### Bacteria-killing assay

Bone marrow-derived monocytes were prepared, adhered, stimulated with IFN-γ and then infected with bacteria as previously described (80). Briefly, nonopsonized *E. coli* DH1 (OD_600_ = 0.6) were added to inflammatory monocyte cultures at a ratio of 20:1. Chambers were spun in an Eppendorf 5810 R centrifuge at 1000 x rpm for 1 min and then transferred to 37°C for 20 min to allow binding and phagocytosis, after which noninternalized bacteria were removed by four washes with PBS containing 5 μg/mL gentamycin (Life Technologies). Cells were further incubated at 37°C for either 20 min or 4 h, in complete medium supplemented with 5 μg/mL gentamycin. Upon washing with PBS, bactericidal activity was assessed via lysis with a solution of 1% Triton X-100 (v/v, in PBS), serial dilutions (1/2000 was optimal), plating on agar, incubation for 18 h at 37°C and counting of CFUs per milliliter. Where indicated, inflammatory monocytes were pretreated for 1 h with the NADPH oxidase inhibitor DPI (10 μM) or with vehicle drug (DMSO 0.1%) before adding bacteria.

### Phagocytosis

Bone marrow-derived monocytes were prepared, adhered and stimulated with IFN-γ as described above. Prior to phagocytosis, zymosan particles and metacyclic promastigotes were washed three times with Hank’s Balanced Salt Solution (HBSS) (Life Technologies), incubated with serum from C5-deficient DBA/2 mice (Charles River Laboratories) for 30 min at 37°C and washed thrice again. To synchronize phagocytosis, cells were fed opsonized zymosan particles or metacyclic promastigotes in cold complete medium without IFN-γ, spun in a Sorvall RT7 centrifuge for 1 min at 1000 rpm and transferred to 37°C to trigger internalization. After 15 min, excess particles or parasites were removed by washing three times with warm medium. Cells were then further incubated in complete medium at 37°C for the indicated times. To monitor phagosome acidification, LysoTracker Red DND-99 reagent (Molecular Probes, diluted 1:2000) was added to cultures 2 h before the end of the indicated time points.

### Light microscopy

To detect superoxide production, inflammatory monocytes fed zymosan particles or infected with metacyclic promastigotes as described above were incubated with 1 mg/mL NitroBlue Tetrazolium (Molecular Probes) for 1 h at 37°C. For parasite survival and proliferation analyses, cells were infected for 2 h with metacyclic promastigotes, washed to remove noninternalized parasites and then incubated for the indicated times. Permanox plastic slides were detached from the 8-well chambers, stained with the Hema 3 Stat Pack (Fisher Scientific) and finally covered with coverslips mounted on Fluoromount-G (SouthernBiotech) prior to sealing with nail polish. A minimum of 100 phagosomes per experimental replicate were examined on a Nikon Eclipse E800 microscope. Images were taken using a Leica DM4000 B microscope equipped with a Leica DFC320 digital camera.

### Confocal immunofluorescence microscopy

After phagocytosis and incubation for the indicated times, inflammatory monocytes were washed with PBS, fixed with 2% paraformaldehyde (PFA; Canemco and Mirvac) for 20 min and then simultaneously blocked and permeabilized in 0.1% Triton X-100, 1% bovine serum albumin, 20% normal goat serum, 6% non-fat dry milk and 50% FBS for 20 min. For the visualization of NADPH oxidase subunit recruitment, cells were sequentially incubated with a solution containing mouse anti-gp91^*phox*^ (BD Transduction, clone 53, diluted 1:70) and rabbit anti-p47^*phox*^ (Upstate, cat. 07-500, diluted 1:100) antibodies for 2 h, followed by a combination of AlexaFluor 488-conjugated goat anti-mouse IgG and AlexaFluor 568-conjugated goat anti-rabbit IgG (Molecular Probes, diluted 1:500) for 30 min, and finally with a mixture of AlexaFluor 647-conjugated rat anti-mouse Ly6C antibody (BioLegend, clone HK1.4; diluted 1:500) and DAPI (Molecular Probes) for 30 min. For phagosome acidification analyses, after fixation and permeabilization, cells were only labelled with DAPI and Ly6C antibody, for 30 min. Cells were washed three times with PBS between each incubation and all steps were performed at room temperature. After staining, Permanox slides were detached from the 8-well chambers and covered with coverslips mounted on Fluoromount-G prior to sealing with nail polish. Fluorescence analyses were performed with a Plan APOCHROMAT 63x oil-immersion DIC 1.4NA objective on a Zeiss LSM780 confocal microscope (Carl Zeiss Microimaging). Images were acquired in plane scanning mode and were minimally and equally processed using Carl Zeiss ZEN software. For each experimental replicate, 100 to 400 phagosomes in Ly6C-positive cells were examined.

### Statistical analysis

Univariate scatter plot graphs, presenting data as means ± SEM, were prepared using the GraphPad Prism software. Statistical significance was assessed using either one-way ANOVA followed by Bonferroni *post-hoc* tests or using a two-tailed, unpaired t-test (** *P* < 0.01, *** *P* < 0.001).

## Acknowledgements

This work was supported by Canadian Institutes of Health Research (CIHR) grant PJT-156416 to AD. AD is the holder of the Tier 1 Canada Research Chair on the Biology of intracellular parasitism. We are grateful to J. Tremblay for assistance with confocal microscopy experiments.

## Author contributions

Conceived and designed the experiments: CM, GAD and AD. Performed the experiments: CM and GAD. Analyzed the data: CM, GAD and AD. Wrote the paper: CM, GAD and AD. All authors discussed the findings and commented on the manuscript.

